# Destabilization of Ran C-terminus promotes GTP loading and occurs in multiple Ran cancer mutations

**DOI:** 10.1101/305177

**Authors:** Yuqing Zhang, Jinhan Zhou, Yuping Tan, Qiao Zhou, Aiping Tong, Xiaofei Shen, Da Jia, Xiaodong Sun, Qingxiang Sun

## Abstract

Ran (<u>Ra</u>s-related <u>n</u>uclear protein) plays several important roles in nucleo-cytoplasmic transport, mitotic spindle formation, nuclear envelope/nuclear pore complex assembly, and other diverse functions in the cytoplasm, as well as in cellular transformation when activated. Unlike other Ras superfamily proteins, Ran contains an auto-inhibitory C-terminal tail, which packs against its G domain and bias Ran towards binding GDP over GTP. The biological importance of this C-terminal tail is not well understood. By disrupting the interaction between the C-terminus and the G domain, we were able to generate Ran mutants that are innately active and potently bind to RanBP1 (<u>R</u>an <u>B</u>inding <u>P</u>rotein 1), nuclear export factor CRM1 and nuclear import factor KPNB1. In contrast to previously reported activated Ran mutants, the C-terminus destabilized mutants are hydrolysis competent in cells, support nuclear transport, and do not form nuclear rim staining. Crystal structures show that one of these C-terminal mutations slightly changes its mode of binding to RanBP1. Finally, a high percentage of Ran C-terminus mutations from cancer patients were found to be destabilizing and hyperactivating, suggesting that Ran C-destabilization might be an unprecedented cellular transformation mechanism in affected cancers. This study also highlights a new drug design strategy towards treating patients with hyperactivated Ras proteins including K-Ras.

## Introduction

Ran (Ras-related nuclear) protein is a member of the Ras superfamily small GTPases. Like other GTPases, Ran switches between GDP (inactive) and GTP (active) bound states. The chromatin-bound guanine nucleotide exchange factor (GEF) RCC1 (Regulator of Chromosome Condensation 1) increases GDP dissociation rate and charges Ran with GTP in the presence of abundant cellular GTP ^1^. On the other hand, the cytoplasm-localized GTPase activating protein (GAP) RanGAP increases the rate of GTP hydrolysis on Ran ^2^. The restricted localization of RCC1 and RanGAP creates a steep Ran-nucleotide gradient: being predominantly GTP-bound in the nucleus and GDP-bound in the cytoplasm ^3^.

Ran is well studied for its role in nucleo-cytoplasmic transport ^4,5^ In the nucleoplasm, RanGTP unloads nuclear localization signal (NLS)-containing-cargo from an importin and forms a complex with the later ^6–8^. Also in the nucleus, Ran, nuclear export signal (NES)-containing-cargo and an exportin form a trimeric nuclear export complex ^9,10^. Either RanGTP-importin or RanGTP-exportin-NES-cargo then transits through the nuclear pore complex (NPC) to the cytoplasm, where RanGTP hydrolysis by RanGAP terminates different Ran complexes with the help of Ran binding protein 1 or 2 (RanBP1 or RanBP2) ^11,12^. RanGDP is recycled back to the nucleus by nuclear transport factor 2 (NTF2) ^13,14^. Besides nuclear transport function in interphase cells, Ran is also critical for mitotic spindle formation, nuclear envelope assembly and NPC assembly during and after mitosis, and diverse other functions in the cytoplasm ^15–17^. Further, Ran hyperactivation is associated with cellular transformation ^18–20^ and the progression of a few cancers ^21–23^. Particularly, Ran is overexpressed in breast cancer and inhibition of Ran activation using anti-RCC1 peptide has demonstrated preferential cytotoxicity in breast cancer cells ^24,25^. Interestingly, its functions in these different processes are connected to its interaction with importins, exportins, RanBP1/2, etc. To investigate the underlying mechanisms, *in vitro* purification of Ran is often an essential step.

While purification of RanGDP is simple ^26^, purification of active Ran (charged with GTP or GTP analogue) is complicated and inefficient ^8,27^. One strategy is to mutate the catalytic residue Q69 to an L (Q69L) in order to slow down the intrinsic GTP hydrolysis rate ^27^. Alternatively, one could use excess of slower hydrolyzing GTP analogue GppNHp to charge Ran in the presence of alkaline phosphatase (AP) ^28,29^. Both methods require an activation protocol which tends to partially inactivate Ran, and the yield of active Ran is estimated to be 30% −80% in the absence and presence of AP respectively ^2,28,29^. Another way of activating Ran is through deletion of its C-terminal 40 residues or only the C-terminus DEDDDL residues ^30,31^. Unlike Ras superfamily proteins (such as Ras, Rab and Arf), Ran contains a unique C-terminal tail that packs against its G-domain ^32^, probably accounting for the tenfold lower affinity for GTP compared with GDP ^27^. However, C-terminal region is also critical for binding of RanBP1 and RanBP2, which are very important effectors of Ran ^11,12,33^. Though Ran is robustly activated after C-terminus deletion ^29^, its interactions with RanBP1/2 are concomitantly abolished, limiting the application of these Ran mutants ^33,34^.

Despite the approaches discussed above, we were unable to generate a highly-active form of Ran required for our study (see discussion). In attempts to search for active Ran which binds to RanBP1/2 and without the usage of expensive materials such as GppNHp and AP-conjugated beads, we designed four mutations to disrupt the interaction between the C-terminus and the G domain. We purified these mutants together with Ran^WT^, Ran^Q69L^, and C-terminus deletion, mutants and compared their activities towards RanBP1, nuclear export factor CRM1, nuclear import factor KPNB1 (also known as importin beta 1), RCC1, RanGAP and NTF2. By X-ray crystallography, we visualized their mode of binding to RanBP1. Further, cellular localization of these mutants and their ability to support nuclear transport in HeLa cells were analyzed. Finally, we discovered several C-terminus destabilizing and hyperactivating (above normal level of activation) Ran cancer mutations, possibly explaining their mechanism of pathogenesis.

## Materials and Methods

### Cloning, protein expression and purification

The human Ran mutants were cloned separately into pET-15b expression vectors incorporating an N-terminal his-tag fusion. Expression of his-Ran was induced by the addition of 0.5 mM isopropyl β-D-1-thiogalactopyranoside (IPTG) in *E. coli* BL21 (DE3) cells, and the culture was grown four hours at 37°C in LB Broth (Miller). Cells were harvested and sonicated in lysis buffer (20 mM Imidazole pH 8.0, 400 mM NaCl, 5 mM MgCl_2_ and 1 mM PMSF). Proteins were purified on a Ni-NTA column and eluted in a buffer containing 300 mM Imidazole pH 8.0, 300 mM NaCl, 5 mM MgCl_2_ and 1 mM beta-mecaptoethanol (BME). This is followed by a Superdex 200 increase gel filtration column on Äkta Pure (GE Healthcare) using gel filtration buffer (20 mM Tris pH 8.0, 100 mM NaCl, 5 mM MgCl_2_, 5 mM BME). Purifications of other proteins were as described previously ^35–39^.

### GTP/GDP quantification

Proteins (500μg, in less than 1ml volume) were briefly denatured by adding 100mM NaOH at room temperature. The denatured samples were added with 10ml of buffer A (10mM Tris pH8.0) to reduce the ionic strength. The samples were loaded onto a Hitrap Q column (GE Healthcare) and eluted with increasing gradient of buffer B (1M NaCl) on Äkta Pure (GE Healthcare). Pure GDP and GTP were eluted at approximately 220mM and 280mM NaCl respectively. The experiments were repeated at least twice to check for consistency.

### Crystallization, data collection, structure solution and refinement

After purification of the complex by Superdex 200 increase gel filtration column, protein complexes were concentrated to 6mg/ml and mixed at 1:1 ratio with crystallization solution containing 18% PEG3350, 200 mM ammonium nitrate, 100 mM Bis·Tris, pH 6.6. 12% (v/v) glycerol was supplemented with crystallization condition as the cryo-protectant. X-ray diffraction data was collected at Shanghai Synchrotron Radiation Facility (SSRF) beamline BL17U1 and BL19U1 ^40^. Coordinates of yCRM1-hRan-yRanBP1 (pdb code: 4HAT) were used as the search model, and refined with rigid body briefly then restrained refinement using the program Refmac5 ^41^. Translation / Libration / Screw (TLS) refinement ^42^ were used in the refinement process. The data collection and refinement statistics are provided in Table S1.

### Pull down assay

To assess different interactions, GST-tagged proteins were immobilized on GSH beads, and an immediate wash step was performed to remove unbound GST tagged proteins. Soluble proteins at indicated concentrations were incubated with the immobilized proteins in a total volume of 1ml for one hour at 4 °C with gentle rotation. After three wash steps, bound proteins were separated by SDS PAGE and visualized by Coomassie Blue staining. Each experiment was repeated at least twice. Pull down buffer contained 20 mM Tris pH 7.5, 200 mM NaCl, 10% glycerol, 2 mM MgCl_2_, 0.001% Triton-X100, and 2 mM DTT if not specified.

### Cell culture, western blot and confocal microscopy

HeLa cells were maintained and analyzed as previously described ^43^. Briefly, cells were maintained in Dulbecco’s modified Eagles medium (Hyclone) supplemented with 10% (v/v) fetal bovine serum (Biological Industries), and transfected with TurboFect transfection reagent (Thermo Scientific). GAPDH (ProteinTech) and mCherry (ProteinTech) antibodies were used at 1:5000 and 1:1000 dilution respectively. Images were acquired by Olympus FV-1000 confocal microscope, and were analyzed using NIH ImageJ and Graphpad software’s.

### *In vitro* nuclear transport using semi-permeabilized cells

The *in vitro* nuclear import assay was slightly modified from reported earlier ^44^. Briefly, 1 μM GST-IBB, 0.5 μM KPNB1, 1 μM NTF2, 1× energy regeneration system ^44^, 0.01% Triton-X100, and 2 μM of different Ran proteins were added to semi-permeabilized HeLa cells and incubated at room temperature for 60 mins. After reaction, the cells were washed, fixed, and visualized by immunostaining with GST antibody. For nuclear export assay, semi-permeabilized HeLa cells were first incubated with 1 μM GST-hRanBP1, 2 μM Ran^WT^ and energy regeneration system for 60 mins to accumulate nuclear GST-hRanBP1. The cells were then incubated with 1 μM of hCRM1, energy regeneration system, 0.01% Triton-X, and 2 μM of different Ran proteins for 30 mins at room temperature with gentle shaking. After reaction, the cells were washed, fixed and visualized by immunostaining with GST antibody. Statistics were based on measurements from at least 30 cells for each sample, and statistical significance was calculated by one-way ANOVA test in Graphpad software.

### Data availability

Structure factor and atomic coordinates were deposited to Protein Data Bank (PDB) with accession codes 5YRO, 5YST, 5YTB and 5YSU.

## Results

### Design and purification of Ran mutants with destabilized C-terminus

C-terminus (a.a. 180-216) of Ran is packed against its G domain and forms numerous interactions. By analyzing RanGDP crystal structure (pdb:3GJ0), we designed four Ran mutations, namely A133D, L182A, M189D and Y197A, that possibly disrupt binding between the C-terminus of Ran and the G domain (Fig. 1A). Considerations were taken to ensure that RanBP1 binding is not impaired with the help of Ran-RanBP1 crystal structure ^8^. Among these residues, A133 and L182 are strictly conserved from Fungi to human (Fig. S1). Except A133, which resides in the G domain, the other three mutations are located in the C-terminus of Ran, and none of those residues are in direct contact with GTP or Mg^2+^. These mutants were predicted to have a dislodged C-terminal tail, and possibly favor GTP binding over GDP binding. Together with Ran^WT^, Ran^Q69L^, Ran^1-179^, and Ran^1-210^, these proteins were purified by Ni-NTA and size exclusion columns without adding GTP or GTP analogue in any purification stage and quickly frozen in −80 °C after being concentrated to 5-10 mg/ml. The purification yields of these C-destabilized (C-des) mutants were comparable to Ran^WT^.

**Figure 1.**
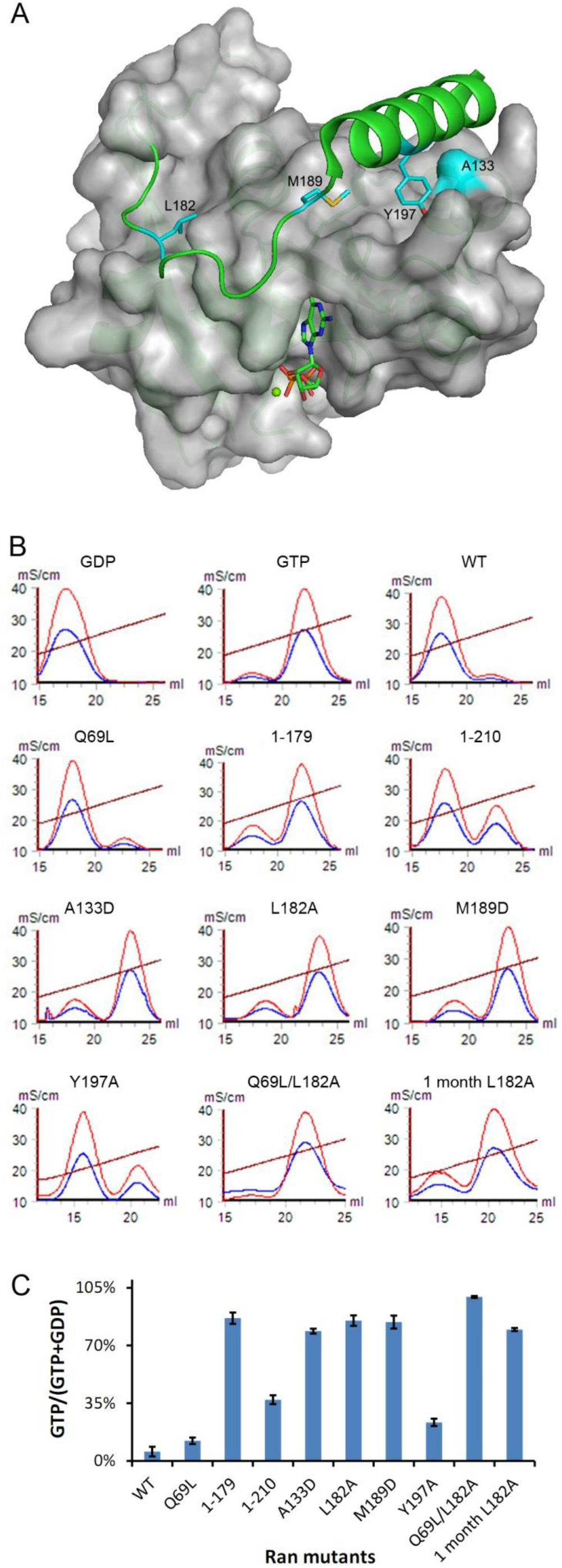
Mutations that destabilize C-terminus of Ran promote GTP loading. A) Ran (3GJ0) is displayed as green cartoon with the G domain covered in partially transparent grey surface. Four loci of mutagenesis (L182, M189, Y197 and A133) are shown as cyan stick representation. The green sphere represents Mg^2+^ ion, and GDP is shown as sticks. B) Q anion-exchange analysis of bound nucleotide in purified Ran proteins. GDP and GTP at 50 μM concentration were used as controls. A260, A280 and conductivity are shown as red, blue and brown lines respectively. C) Percentage of GTP over total bound nucleotide for different Ran proteins. The results shown are an average of two independent purification/quantification experiments. Error bars represent the standard deviation.

### C-des mutants are loaded with higher percentage of GTP

To determine the status of Ran-bound nucleotide, the proteins were denatured by 100 mM NaOH and analyzed by anion exchange Q column (Fig. 1B). GTP and GDP were used as controls to identify GTP and GDP peaks. As expected, Ran^WT^ is merely 5% GTP bound, and without C-terminus, Ran^1-179^ is highly (86%) GTP-bound (Fig. 1B). Though Ran^Q69L^ does not hydrolyze GTP, only 12% of Ran^Q69L^ is GTP-bound. Strikingly, the C-des mutants are significantly more GTP-charged, especially for Ran^A133D^, Ran^L182A^, and Ran^M189D^, ranging from 78%-85% GTP-bound (Fig. 1B). Ran^Y197A^ is loaded with 23% GTP (Fig. 1B).

Though the intrinsic GTP hydrolysis of Ran is slow ^27^, reduced level of bound GTP was observed for Ran^L182A^ after one year of storage at −80 °C (data not shown). To prevent intrinsic GTP hydrolysis, Q69L/L182A double mutant was generated, and its GTP-loading level was analyzed as described above. Surprisingly, Ran^Q69L/L182A^ is loaded with 100% GTP, suggesting that C-des mutation L182A completely switched Ran’s preference to bind GTP over GDP (Fig. 1B). Since time between protein expression in E. coli and GTP% quantification determines the extent of intrinsic hydrolysis, we repeated the quantification experiments with all proteins freshly purified in parallel and found those results reproducible (Figure 1C). In the following experiments we always used the same batch of proteins within a month’s time stored at −80 °C, since one-month-old Ran^L182A^ was similarly GTP-loaded as freshly purified Ran^L182A^ (Fig. 1B,C, 85% versus 81% GTP).

### C-des mutants bind to effector proteins tighter

Next, we analyzed the activity of Ran mutants in binding to different Ran effectors. RanBP1 plays important roles in relieving karyopherin blockage of RanGTP hydrolysis and nuclear export cargo dissociation, displaying a high affinity for RanGTP but not RanGDP. We first tested the binding to GST-tagged RanBP1 by a different concentration of Ran^WT^, Ran^Q69L^, and Ran^L182A^ (Fig. 2A). Unexpectedly, Ran^L182A^ bound to RanBP1 at all concentrations, while Ran^WT^ and Ran^Q69L^ are gradually bound with increasing concentration of Ran, in good agreement with the GTP% observed in Figure 1. When all Ran mutants were tested at the same concentration (1 μM), only the C-destabilized mutants bound strongly to RanBP1, but not Ran^WT^, Ran^Q69L^, Ran^1-179^, or Ran^1-210^ (Fig. 2B). Though Ran^1-179^ was highly GTP-bound, it did not bind to RanBP1 due to lack of C-terminal tail as expected. Negative controls using GST-KPNA2 (importin alpha 1) showed no binding, suggesting that the binding to RanBP1 are specific (Fig. S2).

**Figure 2.**
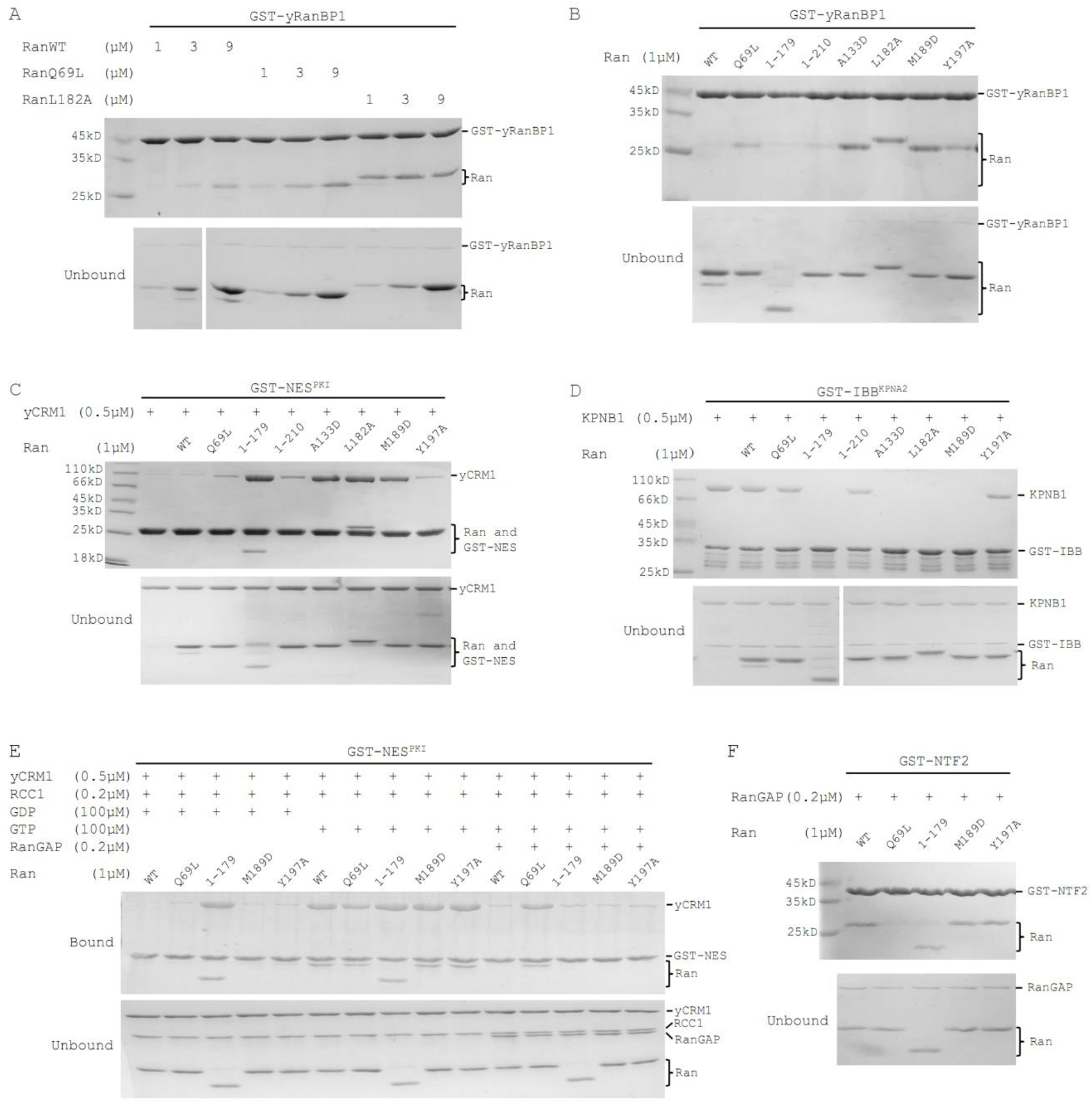
Activity comparison of Ran mutants towards RanBP1, KPNB1, CRM1, RCC1, RanGAP and NTF2. A) GST-tagged yeast (Saccharomyces cerevisiae) RanBP1 (GST-yRanBP1) pull down of Ran^WT^, Ran^Q69L^ or Ran^L182A^ at different concentrations. B) GST-yRanBP1 pull down of Ran and different mutants at fixed concentration (1μM). C) GST-NES^PKI^ pull down of yeast CRM1 (yCRM1) and different Ran proteins. D) GST-IBB^KPNA2^ pull down of KPNB1 in the presence of Ran and mutants. E) GST-NES^PKI^ pull down of yCRM1 and different Ran proteins, in the presence or absence of RCC1, RanGAP, GTP and GDP. F) GST-NTF2 pull down of yCRM1 and different Ran proteins in the presence of RanGAP. RanGAP ensures all Ran proteins are in GDP charged form (Ran^Q69L^ is mainly GDP-charged when purified).

In the nucleus, RanGTP forms a nuclear export complex with CRM1-NES and dissociates KPNB1 cargoes by directly binding to KPNB1. Consistent with GTP% loaded, the C-des mutants were much stronger in forming complex with CRM1-NES or KPNB1, compared with Ran^WT^ (Fig. 2C, D). In addition, GST-NES pull down assay showed no significant activity (CRM1 binding) differences for Ran^L182A^ purified in the presence or absence of GTP (Fig. S3), suggesting that it is unnecessary to add GTP during purification.

### C-des Ran mutants respond to RanGAP and RCC1

To learn whether these Ran proteins respond to RanGAP or RCC1, and whether they bind to NTF2 when in GDP-bound form, we then focused on five representative proteins, Ran^WT^, Ran^Q69L^ (unable to hydrolyze GTP), Ran^1-179^ (86% GTP-bound, but unable to bind RanBP1), Ran^M189D^ (84% GTP-bound), and Ran^Y197A^ (23% GTP-bound). When incubated with either RCC1/GDP or RCC1/GTP, clear differences in amount of bound CRM1 were observed for Ran^WT^, Ran^Q69L^, Ran^M189D^ and Ran^Y197A^, suggesting that these proteins responded to RCC1 activation (Fig. 2E, lane 1-10). It seems that Ran^1-179^ is less sensitive to RCC1 because increasing concentration of RCC1 by tenfold (in the presence of GDP) did abolish CRM1 binding (Fig. S4). As expected, all Ran proteins were sensitive to the addition of RanGAP except Ran^Q69L^, which lacks the catalytic Q69 residue (Fig. 2E, lane 6-15). When these Ran proteins were in the GDP-bound form (by addition of RanGAP), all bound to NTF2 except Ran^Q69L^, since residue Q69 lies in the contact interface ^37^. The results obtained with C-des mutants are consistent with earlier crystal structures which showed that C-terminus of Ran is not involved in binding to RanGAP, RCC1, or NTF2 ^37–39^. In summary, unlike RanQ69L or Ran1-179, C-des mutants RanM189D and RanY197A can be deactivated by GAP-mediated GTP hydrolysis, are sensitive to RCC1 mediated nucleotide exchange, and bind to NTF2 when in GDP-bound form.

### Mode of RanBP1 binding by three C-des mutants

Since C-terminus of Ran is also involved in RanBP1 binding, we crystallized three C-des mutants (Ran^L182A^, Ran^M189D^ and Ran^Y197A^) in complex with RanBP1 and CRM1 in order to examine whether these mutations alter RanBP1 binding ^34^. CRM1 was used because it helped the crystallization process. However, since CRM1 is distant from the mutation sites (Fig. S5), it should not perturb the Ran-RanBP1 binding and is omitted in the following figures to improve clarity. As expected, C-des Ran mutants largely bound to RanBP1 in similar mode (Fig. 3A-C), with all atoms RMSD between 0.3 Å to 0.8 Å. The C-terminus of Ran^L182A^ and Ran^Y197A^ are highly identical as the WT protein (Fig. 3D) ^36^. However, C-terminus of Ran^M189D^ mutant shows significant changes in RanBP1 binding (Fig. 3E, F). In Ran^L182A^ complex structure, M189 is loosely packed on the edge of a hydrophobic pocket in RanBP1 (Fig. 3F). In Ran^M189D^ complex structure, being more hydrophilic, D189 is flipped out towards the solvent, and a previously solvent exposed proline (P191) is inserted into the hydrophobic pocket mentioned above (Fig. 3F). The movement of P191 drags towards the pocket a one-turn helix (A192 to A195), which is originally part of a longer helix (A192 to T206) (Fig. 3E). In addition, the adjacent end of the long helix is shifted about 2 Å from its original position (Fig. 3E arrow). The electron density for C-terminal region of Ran^M189D^ is slightly improved compared to other mutants (Fig. S6), suggesting possibly tighter binding for this mutant. In summary, these structures show that Ran^L182A^ and Ran^Y197A^ bind to RanBP1 similarly, while Ran^M189D^ displays significant changes.

**Figure 3.**
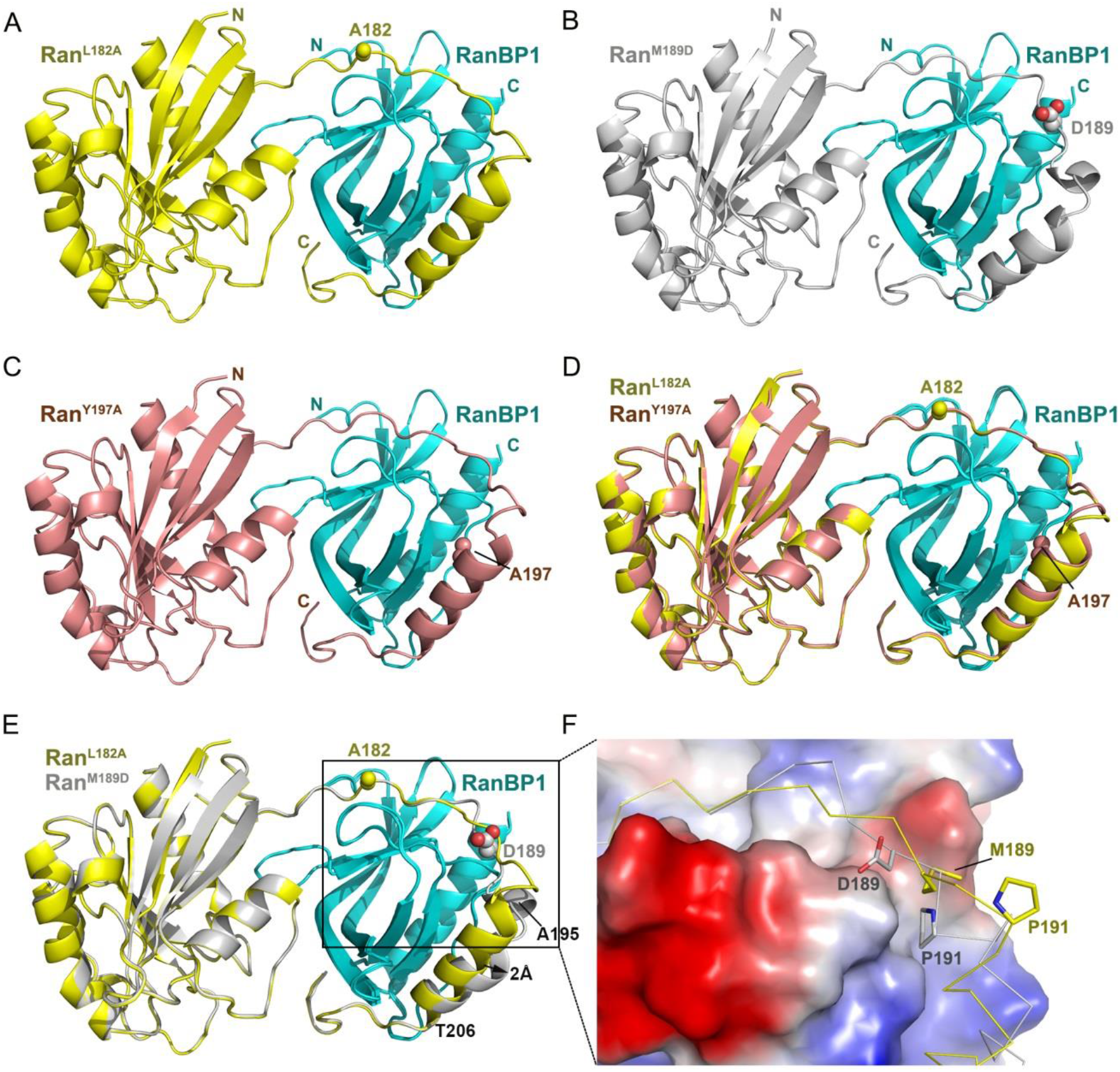
Mode of RanBP1 binding by designed Ran mutants. A, B, C) Crystal structure of Ran^L182A^ (yellow), Ran^M189D^ (grey) and Ran^Y197A^ (salmon) in complex with yeast RanBP1 (cyan) and yeast CRM1. Since CRM1 does not interact with mutated regions, its structures are omitted in these figures to improve clarity. Mutated residues are shown in sphere representation. D) Superimposition of Ran^L182A^-RanBP1 with Ran^Y197A^-RanBP1. E) Superimposition of Ran^L182A^-RanBP1 with Ran^M189D^-RanBP1. F) Zoom in on the boxed region in panel E, displayed with the electrostatic surface map of RanBP1. Residue 189 and 191 are shown in stick representation.

### Activation level and cellular localization of Ran mutants in human cells

To examine whether C-des Ran mutants are activated in human cellular environments, we transfected 293T cells with plasmids encoding mCherry-tagged Ran proteins, lysed the cells, incubated the lysate with immobilized GST-hRanBP1, and blotted Ran using mCherry antibody (Fig. 4A). In contrast to Ran^WT^, which did not bind to RanBP1, Ran^Q69L^, Ran^Y197A^, and Ran^M189D^ were all bound and likely activated in 293T cells. Interestingly, Ran^M189D^ was much more activated than Ran^Q69L^, possibly due to improved binding to RanBP1, as shown by crystal structures (Fig. S6). Alternatively, this observation could argue that preference for GTP binding is more critical than hydrolysis capability in determining Ran’s cellular nucleotide state. Though Ran^1-179^ did not bind to RanBP1, it does not mean that Ran^1-179^ is not charged with GTP because RanGTP^1-179^ does not bind to RanBP1, as shown earlier (Fig. 2B).

**Figure 4.**
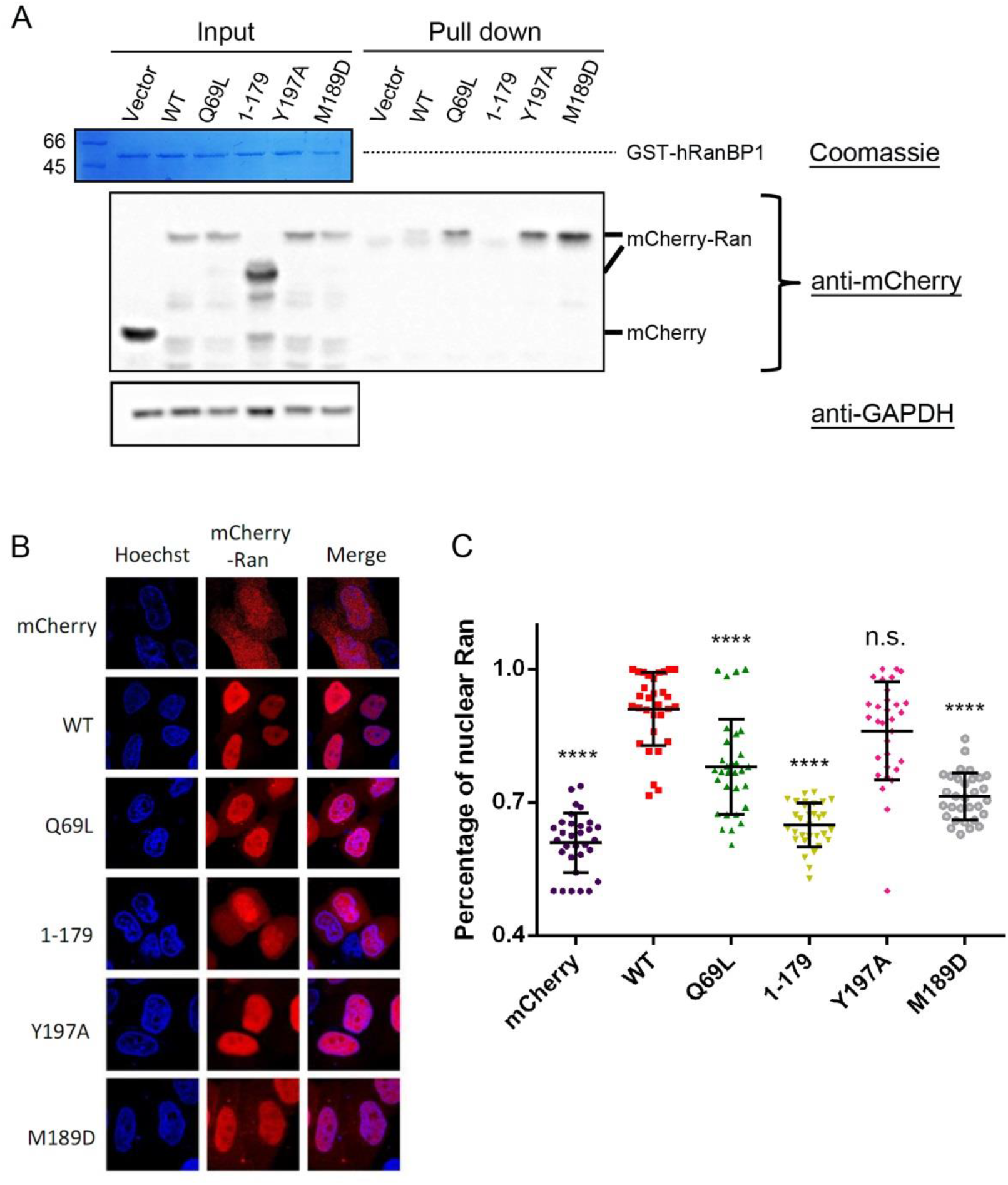
GTP-loading level and cellular localization of Ran mutants in human cells. A) GST-hRanBP1 pull down of mCherry-Ran constructs transiently expressed in 293T cells. Ran proteins were stained with mCherry antibody. B) Intracellular localization of mCherry tagged Ran proteins in transfected HeLa cells. C) Quantification and statistical analysis of localization in 4B. Percentage of nuclear Ran for each cell is calculated as Ran nuclear intensity divided by total cellular intensity. Error bars represent standard deviation of each set of data containing measurements from at least 30 cells. Statistical significance was calculated between Ran^WT^ and other samples. **** denotes p<0.0001; *** denotes 0.0001<p<0.001.

We then studied how these C-destabilized Ran mutants are localized in cells using mCherry-Ran plasmids transfected into HeLa cells. Ran^WT^ and Ran^Y197A^ displayed similar (91% vs. 86%, n.s.) nuclear localization level (Fig. 4B, C). However, a significant fraction of Ran^Q69L^, Ran^1-179^ and Ran^M189D^ were localized in the cytoplasm (78%, 64% and 71% nuclear respectively). In addition, nuclear rim staining was observed as reported for Ran^Q69L^ and occasionally for Ran^1-179^, since they are not hydrolysable in cells and hence stuck on the NPC ^29,45^. Nuclear rim staining was not observed for C-des mutants Ran^M189D^ and Ran^Y197A^, possibly because of hydrolysis competency, as shown in Figure 2E. In summary, C-des Ran mutants are potently activated in eukaryotic cellular environment and tend to localize to the cytoplasm.

### C-des mutants support nuclear transport

Using purified proteins and semi-permeabilized cells, we assessed whether these Ran proteins facilitate nuclear transport of cargoes. Compared with ‘no Ran’ sample (mean:0.02), nuclear import of GST-IBB (<u>I</u>mportin <u>B</u>eta <u>B</u>inding domain of importin alpha) in the presence of nuclear import factor KPNB1 is stimulated by Ran^WT^ (0.53), partially by Ran^Q69L^ (0.22), but not by Ran^1-179^ (0.07) (Fig. 5A), consistent with earlier reports ^29,46^. Ran^Y197A^ (0.47) and Ran^M189D^ (0.42) are similar as Ran^WT^ in promoting nuclear import (Fig. 5A, B). On the other hand, nuclear export of GST-hRanBP1 (which contains a NES) in the presence of nuclear export factor CRM1 was promoted by Ran^WT^ (0.18, the less the number, the higher the export activity), Ran^Y197A^ (0.24), and partially by Ran^M189D^ (0.42) (Fig. 5C, D). Ran^Q69L^ (0.60) and ‘no Ran’ (0.72) samples are statistically insignificant. Interestingly, nuclear cargo intensity of Ran^1-179^ (0.90) samples are significantly (P<0.01) higher than that of ‘no Ran’. We are unclear of the reason, but Ran^1-179^ may act through inhibiting passive diffusion speed of nuclear pores. These results support the notion that hydrolysis competency is more important than being constantly activated for Ran to facilitate nuclear transport. In summary, C-des mutants support nuclear transport in contrast to previously-reported Ran mutants.

**Figure 5.**
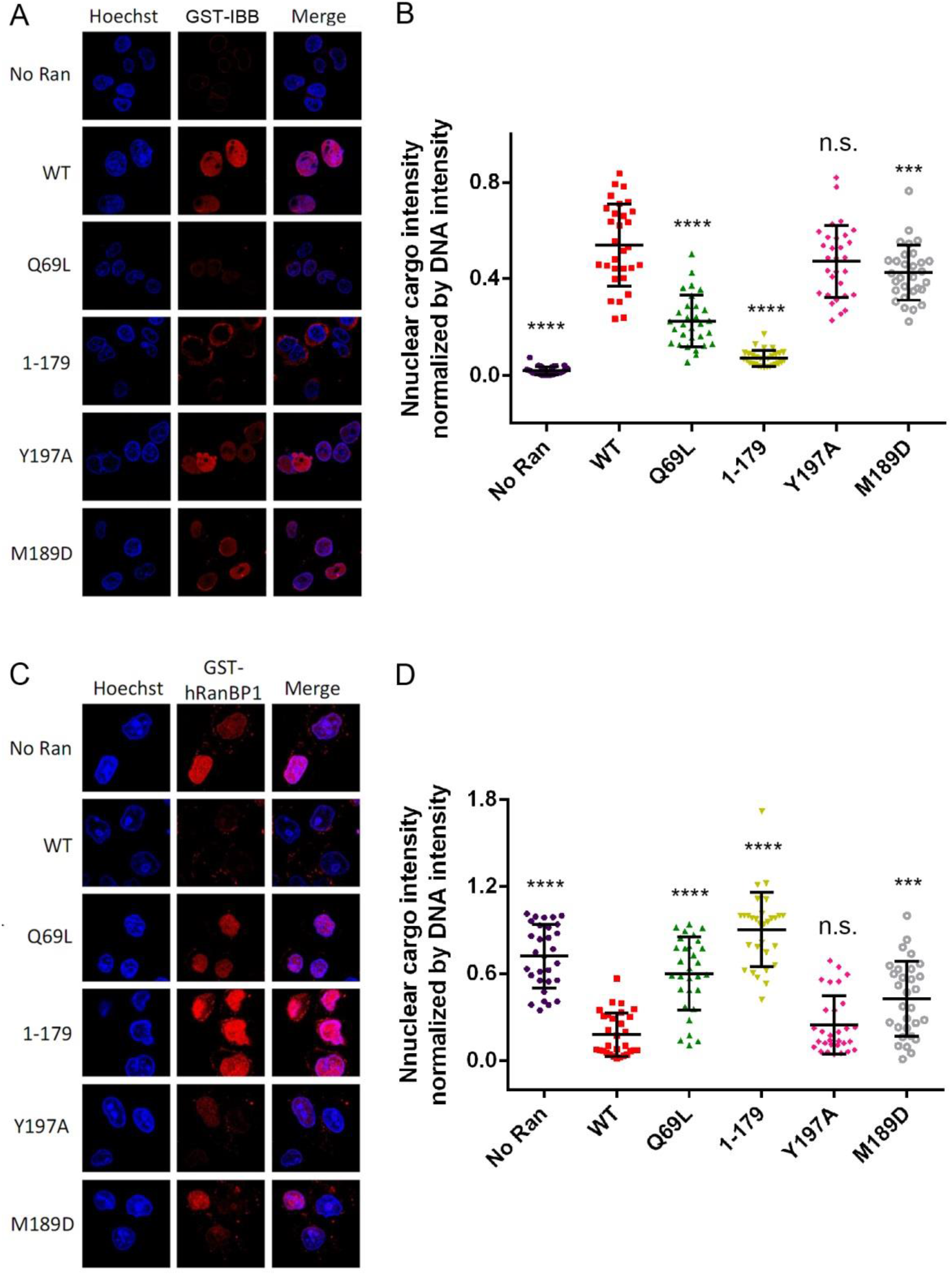
Assessment of nuclear import and export stimulation by Ran mutants in semi-permeabilized HeLa cells. Cells after nuclear import (A) and export (C) of GST-cargoes in the presence of Ran and different mutants were stained with anti-GST antibody (red). GST-tagged importin beta binding domain of importin alpha 1 (GST-IBB) was used as an import cargo. GST-tagged human RanBP1 (GST-hRanBP1) which contains an NES was used as an export cargo. Quantification and statistical analysis of nuclear import or export are shown in B and D respectively. Level of nuclear transport is assessed by nuclear cargo intensity (normalized with DNA intensity). Error bars represent standard deviation of each set of data containing measurements from at least 30 cells. Statistical significance was calculated between Ran^WT^ and other samples.

### Ran cancer mutations may function through impairing auto-inhibition of C-terminus

Next, we conducted a database search on COSMIC and cBioportal servers to look for C-terminus destabilized hyperactive Ran from patient tumor samples. Mutations within residue range 177-187 and H30Y were selected and studied since these mutations are not in direct contact with GTP or Mg^2+^ and might perturb auto-inhibition of the C-terminus (Fig. 6A). For instance, A183 is inserted into a small pocket on G domain, and switching to the bulkier residue T could abolish this interaction and release C-terminus of Ran (Fig. 6A). In addition, this somatic mutation was found in colon adenocarcinoma and was predicted to be pathogenic by FATHMM, with a score of 0.96, through Androgen Receptor Signaling Pathway ^47^. H30Y (kidney and liver), V177A (colon), M179I (Endometrial), and P184S (skin) were also predicted to be pathogenic by FATHMM.

**Figure 6.**
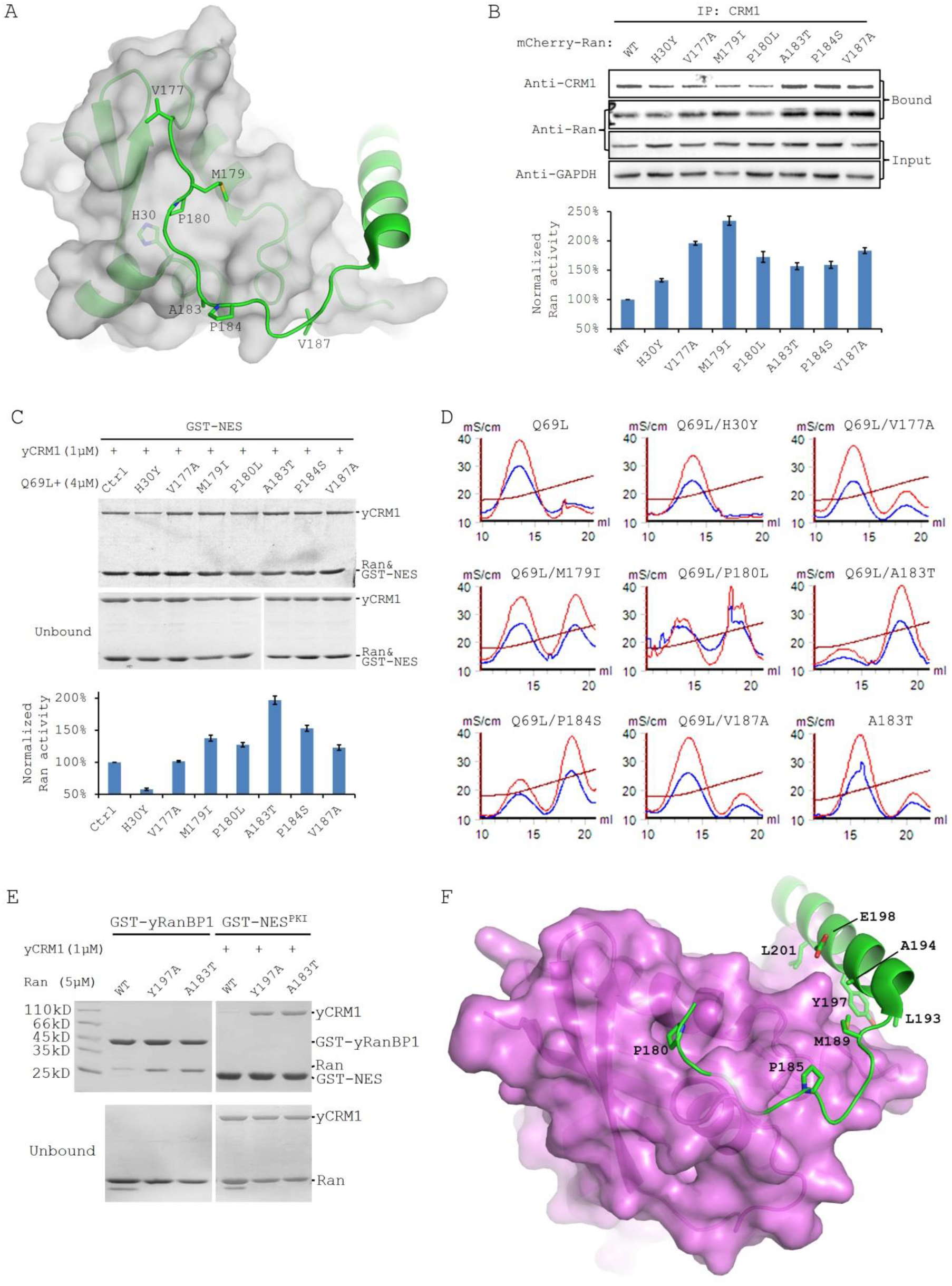
Multiple Ran mutations naturally occurring in human cancers are C-destabilizing and hyperactivating. A) Location of mutated residues (stick) in cancers on RanGDP crystal structure (3GJ0). Ran is displayed as green cartoon with G domain covered in partially transparent grey surface. B) CRM1 immunoprecipitation in 293T cells transiently expressing mCherry-Ran mutants. Bottom panel shows the quantification of Ran activity normalized by CRM1 intensity (gel loading control). C) GST-NES^PKI^ pull down of CRM1 in the presence of Ran^Q69L^ double mutants. Ctrl sample is Ran^Q69L^ single mutant. Bottom panel shows activity of Ran calculated by normalizing CRM1 intensity with GST-NES band intensity. Pull down were repeated twice and checked for consistency. D) GTP% quantification by Q column analysis showed increased level of bound GTP for cancer derived Q69L double mutants. Single mutant Ran^A183T^ was also analyzed and is charged with 23% of GTP. E) Ran^A183T^ displays similar activation level as Ran^Y197A^ in binding to RanBP1 and CRM1. F) Superimposition of RanGDP (pdb:5bxq) onto K-RasGDP (pdb:5W22). KRas is shown as magenta surface. RanGDP C-terminus (green) residues which favorably interact with K-Ras are shown as sticks.

In order to test whether these mutants could alter Ran activity, we transfected each mutant into 293T cells and tested their interactions with CRM1. Strikingly, CRM1 immunoprecipitated 30% – 130% more Ran mutants relative to Ran^WT^, suggesting mild activation of those mutants (Fig. 6B). In order to analyze the level of activation in vitro, we engineered those mutations on top of Ran^Q69L^ mutant and purified those double mutants, including Ran^Q69L^ single mutant in *E.coli*. The purpose of designing Ran^Q69L^ double mutants is to minimize the influence of intrinsic hydrolysis during purification. Pull down assay using *E.coli* expressed proteins showed that except Ran^Q69L/H30Y^ and Ran^Q69L/V177A^, the other mutants are 20%-100% more active than Ran^Q69L^ (Fig. 6C). We further analyzed the GTP% bound by Q column to precisely quantify the level of activation. Clearly, 10-70% increased level of bound GTP was observed for all double mutants except Ran^Q69L/H30Y^ and Ran^Q69L/V187A^ (Fig. 6D, S7). Combining three different approaches above, four mutations (M179I, P180L, A183T, P184S) are constantly shown to be hyperactivating. Since A183T is most hyperactivating *in vitro*, we further generated Ran^A183T^ single mutant and found that it was charged with 23% of GTP, similar as Ran^Y197A^ (Fig. 1B, 6D). Indeed, Ran^A183T^ displayed a similar level of binding to RanBP1, CRM1 and KPNB1 as Ran^Y197A^ by pull down (Fig. 6E, S8), suggesting all previous studies on Ran^Y197A^ possibly apply to Ran^A183T^. In summary, at least four out of seven tested Ran cancer mutations are C-destabilizing and mildly hyperactivating, suggesting that Ran hyperactivation through impaired auto-inhibition might be a novel pathogenic mechanism in various cancers.

## Discussion

### Advantages of the designed C-des Ran mutants

It turns out that C-terminus of Ran is very sensitive to mutation since all designed mutations increase Ran’s activity significantly. These C-des mutants are charged with higher level of GTP, and bind to effector proteins more tightly when expressed and purified in *E.coli* or lysed 293T cells. The designed C-des mutants have the following advantages simultaneously: 1) C-des mutations enable effortless purification of highly GTP-bound Ran. *In vitro* generation of highly GTP-bound Ran is hard since Ran tends to bind GDP over GTP. Especially, the GTP that we bought contains 10% – 40% of GDP (Fig. 1B, GTP from Sigma). Plus the fact that Ran is always purified with more than 90% of GDP, our initial attempts to generate GTP-loaded Ran by GTP-charging failed miserably. C-des mutation completely reverses Ran’s preference for nucleotide and allows easy purification of highly GTP-bound Ran. Especially, we showed that Ran^Q69L/L182A^ double mutant purified from *E. coli* is charged with 100% GTP. 2) The purification process requires neither GTP or GTP analogues, nor the steps of GTP-charging to activate Ran. This greatly reduces the time and cost of purification, and it also prevents the fractional denaturation of Ran during activation. 3) The C-des Ran mutants are able to bind to RanBP1 and RanBP2. This is important because these proteins are essential effectors of RanGTP, playing critical roles in nuclear transport. For example, the formation of CRM1-Ran-RanBP1 complex is possible with C-des mutants, but not with Ran^1-179^ or Ran^1-210^. 4) The C-des mutants are hydrolysis competent *in vitro* and in cells. In contrast to previously reported hydrolysis-incompetent mutants Ran^Q69L^ and Ran^1-179^, C-des mutants do not show nuclear rim staining and support nuclear transport, possibly due to their competency in GTP hydrolysis. Ran^1-179^GTP is not hydrolysable in cells because of incompetency to bind to RanBD (due to lack of C terminus), therefore trapped in importins/exportins proteins ^11,12^.

### Applications of designed C-des mutants

Because of those properties, the C-des Ran mutants or the double mutant Ran^Q69L/L182A^ could be applied in various ways not limited to the examples listed below. In this study, we showed that the C-des mutants are useful to generate protein crystal structures. In pull down experiments, C-des mutant Ran^L182A^ could not only reduce the amount of proteins used, but it could also allow better concentration determination of RanGTP.

In addition, the C-des mutants could be used in various cellular studies because they are hydrolysis competent and do not form artifacts such as nuclear rim staining. Furthermore, it might be possible to design a Ran mutant that constantly binds to GDP by further stabilizing the interaction of C-terminus with the G domain. Such mutants should be useful for cellular imaging studies ^45^.

Ran^Q69L/L182A^ enables one to use accurate protein concentration in an experiment and rules out the contamination of RanGDP. For example, it is necessary to know the exact protein concentrations when doing Isothermal Titration Calorimetry (ITC). Furthermore, RanGDP is shown to bind NTF2, zinc fingers and several other proteins weakly such as KPNB1 and RanBP1. When having contaminating RanGDP is undesirable, using Ran^Q69L/L182A^ double mutant could effectively resolve the problem.

### Ran hyperactivation and cellular transformation

After success with our C-des design, we reason that similar mutations might exist naturally in human cancers, since activation of Ran is reported to be cell transforming and Ran overexpression is observed in different cancers ^18,19,23,24,48^. Albeit relatively weak in level of activation, a high percentage of patient-derived Ran mutants tested are hyperactivated both in cells and *in vitro* demonstrated by immunoprecipitation or pull down. Especially, we showed that the ratios of bound GTP for six double mutants were significantly higher compared to Ran^Q69L^ through Q-column analysis (Fig. S7). Considering that only a small fraction of Ran mutations were tested, there possibly exist more C-destabilizing Ran mutations that promote GTP-loading in cancer patients. Interestingly, highly activated Ran mutations (such as L182A) were not found in cancer patients, likely because that such mutants disrupt cellular RanGTP gradient and inhibit nuclear transport (Fig. 4B, 5), which should be harmful for cancer cells ^49^. Among those mutations tested, Ran^A183T^ from colon cancer patients is similarly hyperactivated as Ran^Y197A^, assessed by GTP% loading and pull downs. Ran^A183T^ and other mildly hyperactivated Ran cancer mutants probably function similarly to Ran^Y197A^ in cells, being predominately localized in the nucleus and supporting nuclear transport. Considering the prevalence of hyperactivation among tested Ran cancer mutations, we believe that C-des mutations should play a role in affected cancers. How C-des mutations contribute to tumorigenesis or cancer progression warrants further studies.

### A possible anti-Ras drug design strategy

Though Ras hyperactivating mutations are frequently observed in different cancer patients, the design of Ras inhibitor has been difficult due to lack of an apparent drug binding pocket on Ras ^50^. Superimposition of RanGDP onto K-RasGDP shows reasonable surface complementarity between C-terminus of Ran and K-RasGDP, with a few C-terminal Ran residues perfectly docked into small cavities on K-Ras (Fig. 6F). It might be possible to design a peptide or small molecule analogous to C-terminal tail of Ran, which loops around G domain of Ras and locks it in its GDP state, to shut down this erroneously hyperactivated oncogene, as a strategy of anti-cancer treatment.

## Conclusion

We designed four C-terminus destabilized Ran mutants that showed higher affinity for GTP, compared to GDP, and thus obtained a high percentage of GTP-bound Ran, even when purified without adding any GTP or GTP analogue, or without performing the previously necessary GTP-charging steps. Pull down assays show that these mutants potently bind to effector proteins and respond to RanGAP or RCC1 mediated GTP hydrolysis or nucleotide exchange. In contrast to Ran^Q69L^ and Ran^1-179^, C-des mutants do not form nuclear rim staining and are able to support nuclear transport, possibly because of hydrolysis competency in cells. Crystal structures show that these mutations bind to RanBP1 similarly, except Ran^M189D^. Finally, from cancer mutation databases we discovered several Ran C-terminal mutations that promote GTP binding through destabilization of C-terminus, providing a possible cellular transformation mechanism in affected cancer.

## Acknowledgements

We thank Dr. Chong Chen and Dr. Rui Bao from Sichuan University for helpful discussions. This research is supported by Natural Science Foundation of China (NSFC) grants (#80502629, #31671477, #81770580). We thank the staffs from BL17U1/BL18U/BL19U1 beamlines of Shanghai Synchrotron Radiation facility (SSRF) for assistance during data collection.

## Author Contributions

Q.Sun conceived the project. Y.Zhang, J.Zhou and Q.Sun performed biochemical, biophysical and cellular works. Q.Sun, D.Jia, X.Sun, and X.Shen supervised the project. Y.Tan prepared several proteins used in this manuscript. Q.Sun prepared the manuscript. Q.Zhou and A.Tong critically revised the manuscript. The authors declare no conflict of interest.

## Supplementary table

**Table 1.**
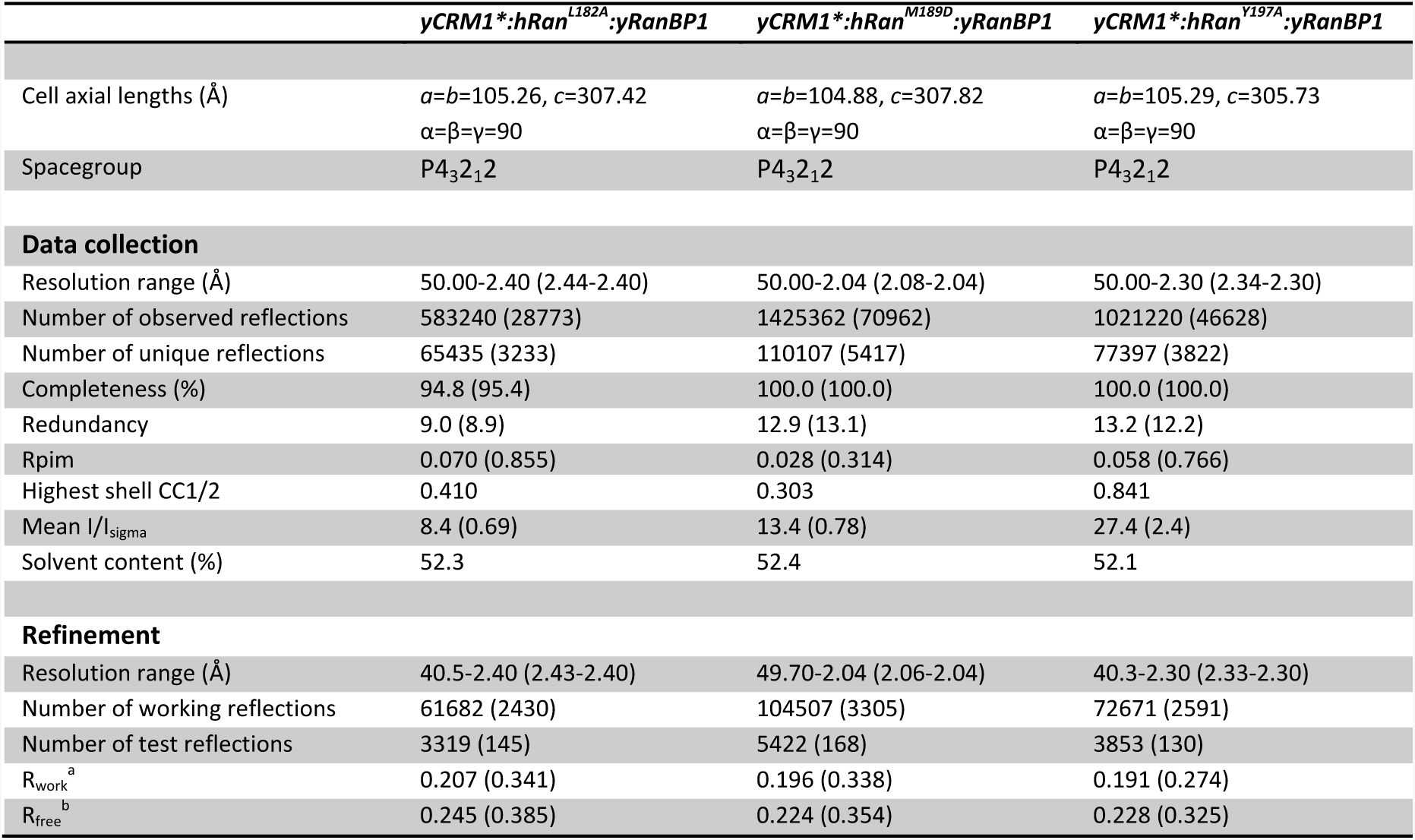
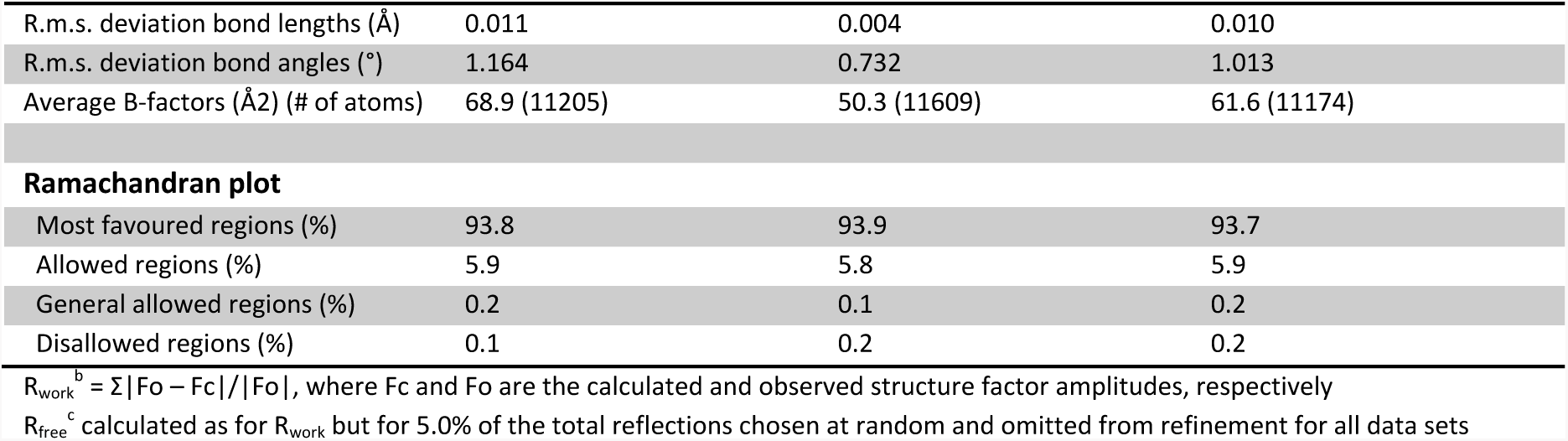
Crystal data collection and refinement statistics.

## Supplementary Figures and Legends

**Figure S1.**
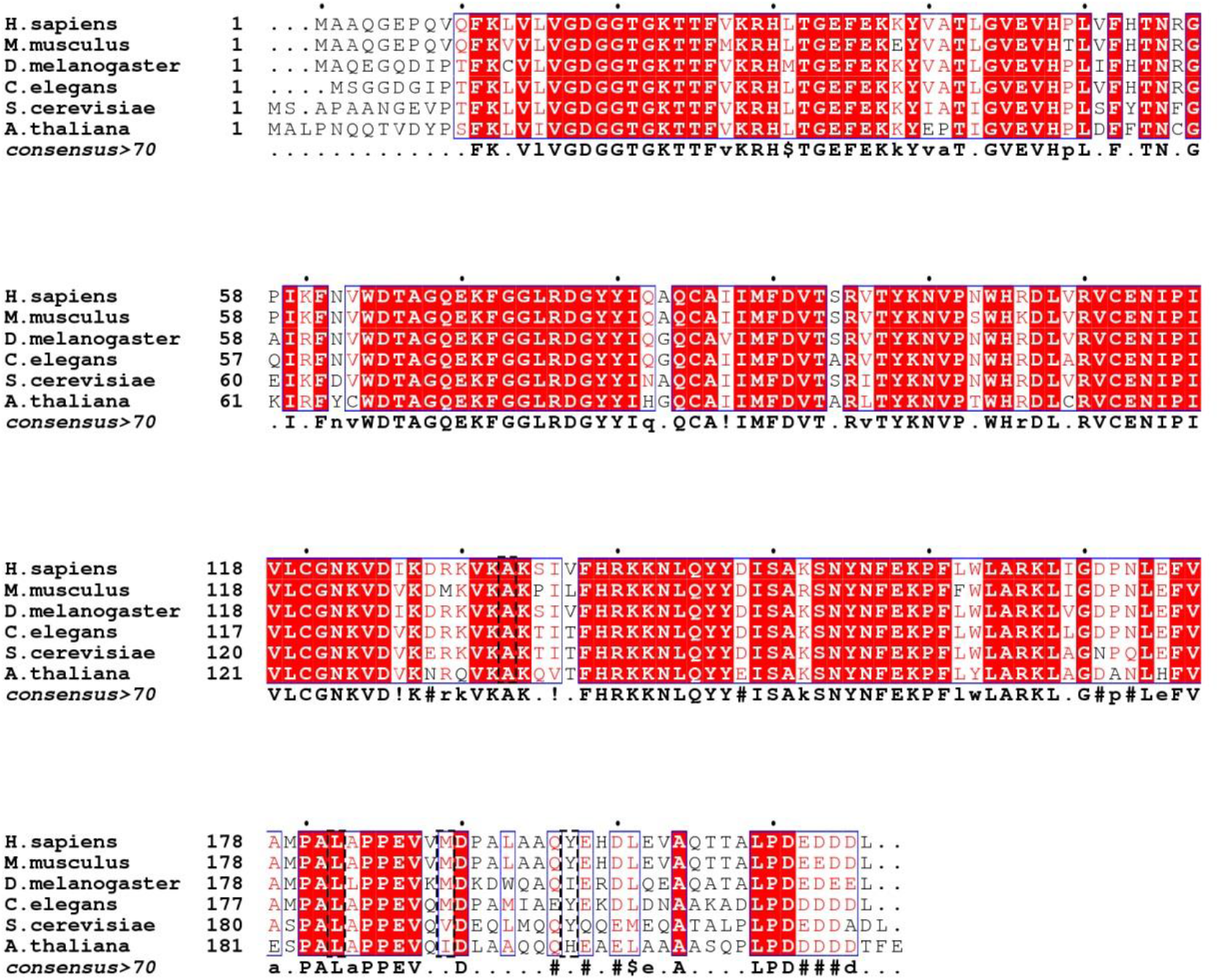
Multiple sequence alignment of Ran from distant species with consensus displayed at the bottom. Mutated residues are boxed.

**Figure S2.**
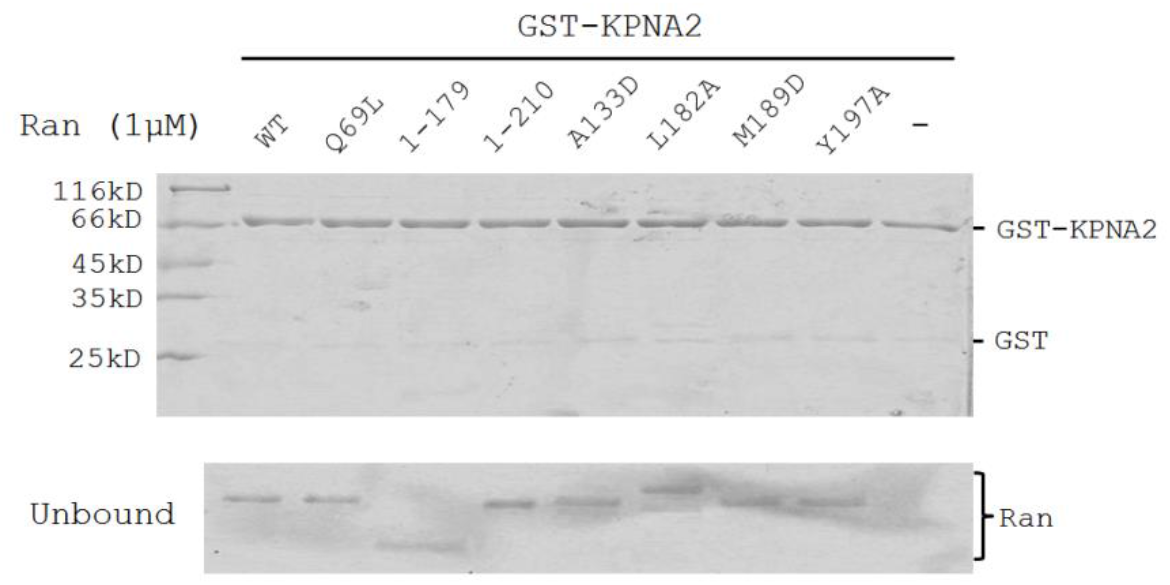
GST-KPNA2 (importin alpha 1) control pull down to show that the binding to RanBP1 is specific. Since Ran and GST are similar in size on SDS-PAGE, GST-KPNA2 (importin alpha 1) rather than GST was used in this pull down.

**Figure S3.**
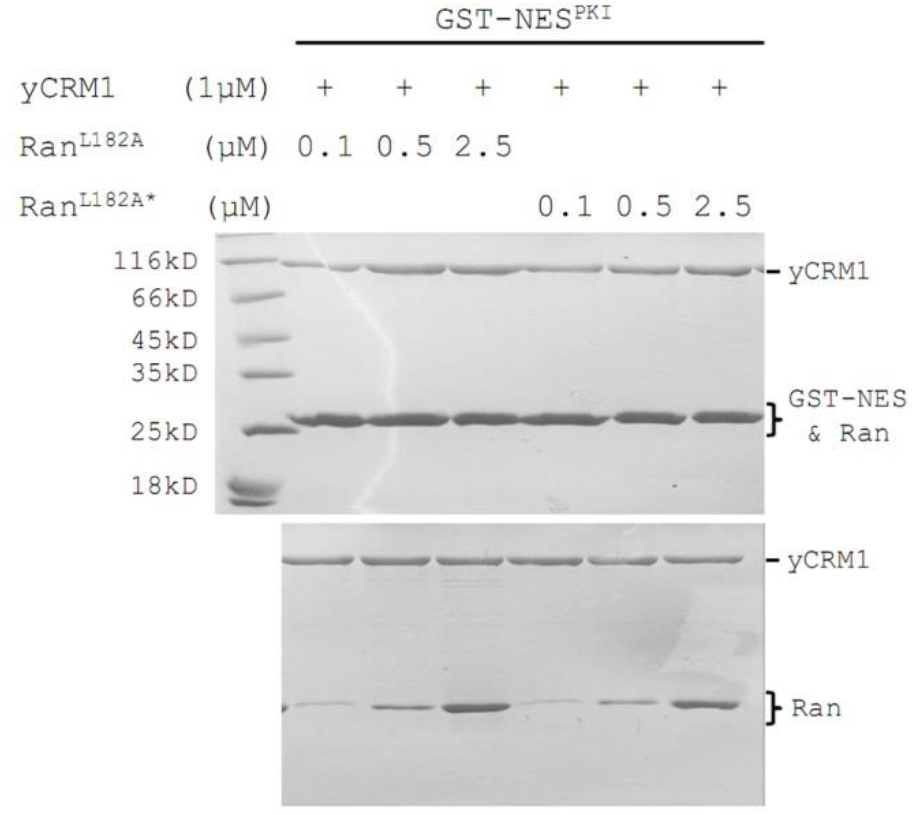
GST-NES pull down of yCRM1 showed no significant activity difference for Ran^L182A^ purified in the presence or absence of 1 mM GTP. Ran^L182A^ purified in the absence of GTP is denoted Ran^L182A*^.

**Figure S4.**
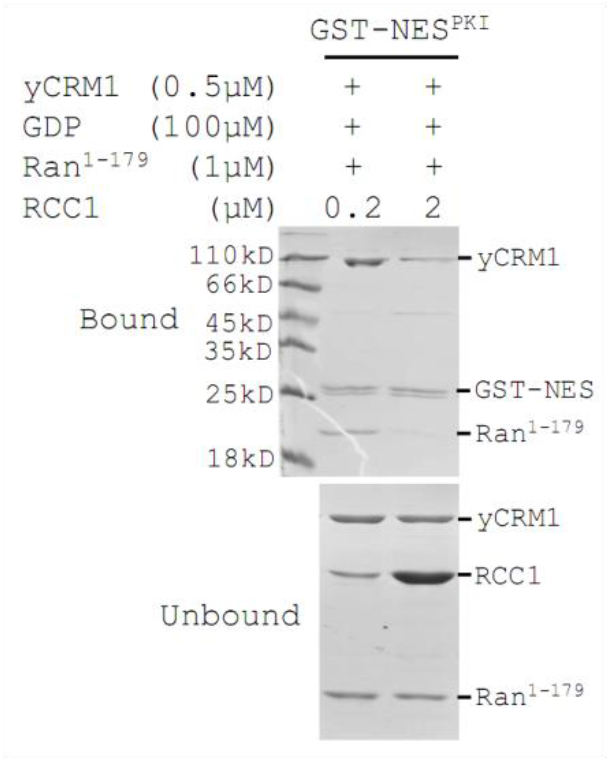
GST-NES pull down of yCRM1 and Ran^1-179^ in the presence of GDP and different concentration of RCC1. yCRM1 and Ran^1-179^ were not bound when concentration of RCC1 is increased to 2 μM.

**Figure S5.**
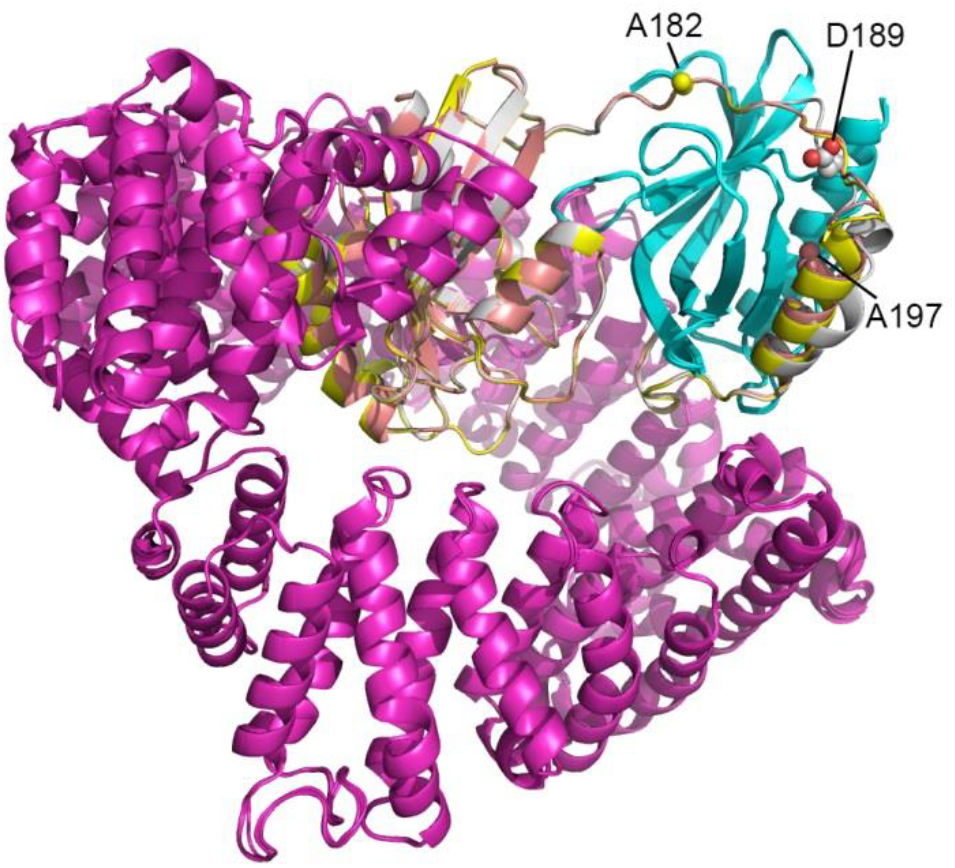
Superimposition of three Ran-RanBP1-CRM1 crystal structures obtained in this study to show that CRM1 is not in contact with mutated Ran residues. CRM1 is shown in magenta color; RanBP1 is shown in cyan color; Ran is shown in yellow (L182A), grey (M189D) and salmon (Y197A) color respectively. Mutated residues are shown in sphere representation.

**Figure S6.**
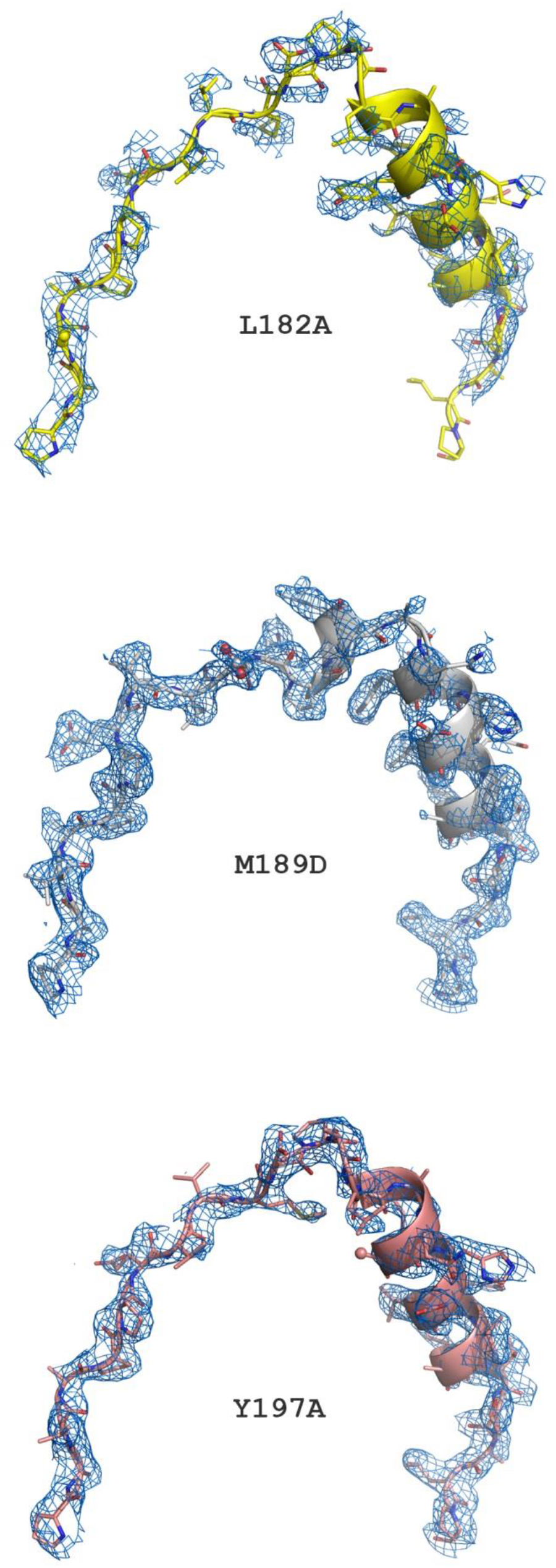
2F_o_-F_c_ omit maps (blue mesh) of Ran C-terminus contoured at 1σ level. Ran is shown as cartoon and stick representation, with the mutated residues shown in sphere. Electron density of Ran^M189D^ is slightly improved, due to minor change in binding RanBP1 after mutation.

**Figure S7.**
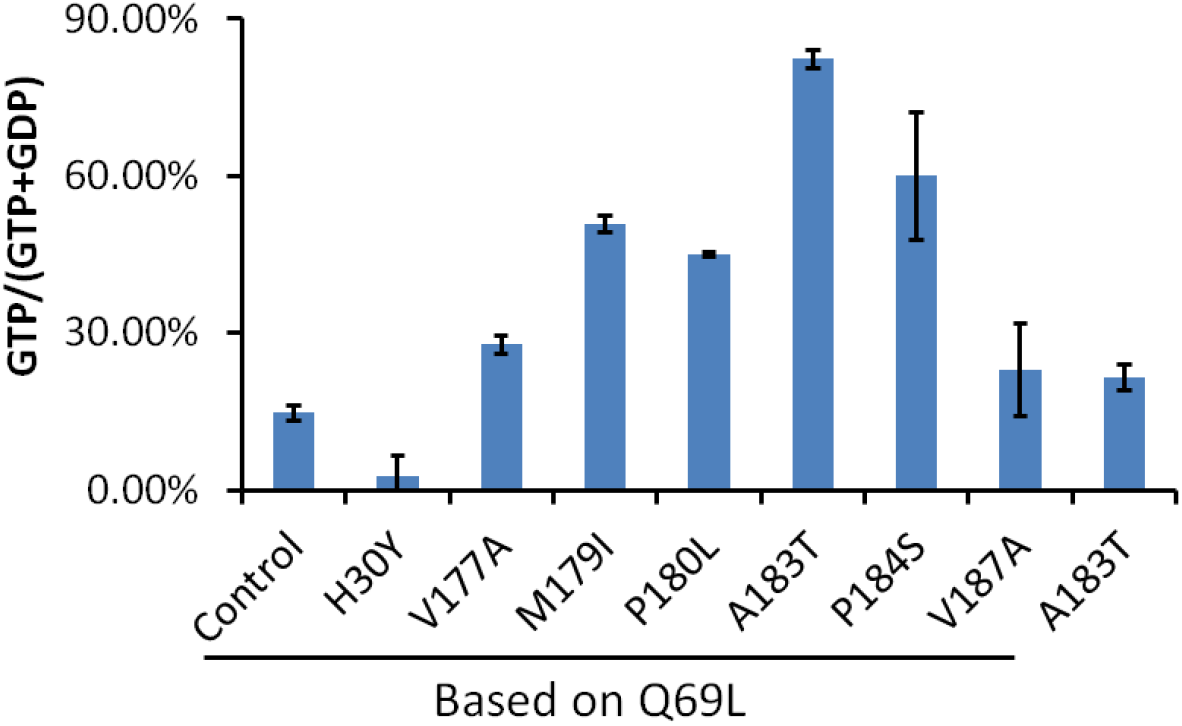
Percentage of GTP over total bound nucleotide for different Ran proteins analyzed by Q anion-exchange column. The results shown are an average of two independent purification/quantification experiments. Error bars represent the standard deviation.

**Figure S8.**
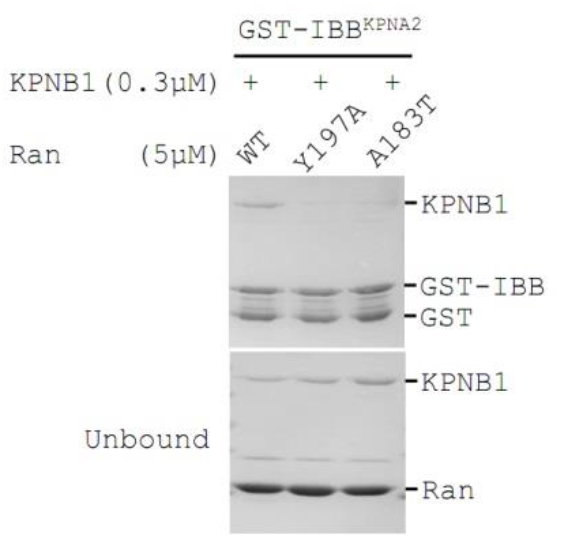
GST-IBB^KPNA2^ pull down of KPNB1 in the presence of Ran^WT^, Ran^Y197A^ and Ran^A183T^.

